# Searching sequence databases for functional homologs using profile HMMs: how to set bit score thresholds?

**DOI:** 10.1101/2021.06.24.449764

**Authors:** Jaya Srivastava, Ritu Hembrom, Ankita Kumawat, Petety V. Balaji

## Abstract

**Motivation:** UniProt and BFD databases together have 2.5 billion protein sequences. A large majority of these proteins have been electronically annotated. Automated annotation pipelines, vis-à-vis manual curation, have the advantage of scale and speed but are fraught with relatively higher error rates. This is because sequence homology does not necessarily translate to functional homology, molecular function specification is hierarchic and not all functional families have the same amount of experimental data that one can exploit for annotation. Consequently, customization of annotation workflow is inevitable to minimize annotation errors.

**Results:** We discuss possible ways of customizing the search of sequence databases for functional homologs using profile HMMs. Choosing an optimal bit score threshold is a critical step in the application of HMMs, which is illustrated using four Case Studies; the single domain nucleotide sugar 6-dehydrogenase and lysozyme-C families, and SH3 and GT-A domains which are typically found as a part of multi-domain proteins. We also discuss the limitations of using profile HMMs for functional annotation and suggests some possible ways to partially overcome such limitations.

**Supplementary information:** Supplementary_material containing Figures S1-S7 and Tables S1 and S2

Supplementary_dataset.xlsx

## INTRODUCTION

<u>H</u>idden <u>M</u>arkov <u>M</u>odel (HMM) is a statistical modelling approach that has gained success across diverse fields (Rabiner et al. 1983; Lander and Green 1987; Churchill 1989; Yang et al. 1994). Profile HMMs have been used for the identification of sequence homologs (Krogh, Brown, et al. 1994), prediction of coding regions (Krogh, Mian, et al. 1994), fold recognition (White et al. 1994; Hubbard and Park 1995), etc. Profile HMMs were introduced in sequence analysis as models to capture sequence variability in all homologous positions of a sequence family of proteins (Krogh, Brown, et al. 1994). Profile HMMs outperform pairwise BLAST in detecting remote homologs (Park et al. 1998) because BLAST does not have a mechanism to consider position specific sequence variations among homologs. Profile HMMs, like PSI-BLAST, use position-specific scores but in addition, they penalize gaps in conserved regions much more than those in variable regions (Durbin et al. 1998; Stojmirović et al. 2008). Consequently, profile HMM databases such as Pfam (Punta et al. 2012) and SUPERFAMILY (Gough et al. 2001) are being used for functional annotation of new protein sequences.

The effectiveness of an algorithm is critically dependent on the values one sets for its parameters. For algorithms designed to identify (remote) homologs, suggested default values are tuned to make the search sensitive, rather than specific. HMMER developers suggest E-value = 10 as the default threshold for *hmmsearch* (Finn et al. 2011) i.e., if one were to search a database of size *S*, even sequences which score *t* bits and E-value 10 will be output as hits. Clearly, this threshold leads to (very) low specificity and hence is unsuitable for sequence-similarity based functional annotation. Thus, a critical step in implementing profile HMMs for detecting functional homologs is the identification of an optimal E-value or bit-score threshold. Choosing an optimal threshold is not straightforward primarily because (i) sequence homology does not necessarily translate to functional homology and (ii) there are not adequate number of experimentally characterized sequences in every family of functional homologs. With an automated process for setting bit score thresholds, optimality cannot be guaranteed since every family of functional homologs is unique. Instead, customizing a workflow taking into consideration the nature and amount of available experimental data can lead to higher prediction accuracies. Herein, we illustrate this approach using two functional families and two domains as Case Studies. It will not be out of place here to mention that we have a catch-22 situation on hand since limitations on the resources for performing experiments mandate the use of prediction methods and validation by experiments is the only way to know the veracity of (large scale) predictions.

## METHODS

Databases and web servers used in this study are listed in Table S1. Input dataset for building profile HMMs consisted of experimentally characterized proteins and reviewed sequences from Swiss-Prot (Figure 1). This type of dataset is henceforth referred to as Exp_dataset. Exp_datasets for Case Studies 1 and 2 were obtained from literature (searched using Google Scholar) and Swiss-Prot, followed by manually verifying that activities have been tested by product characterization, direct enzyme assay, or gene disruption or complementation. The keywords GlcNAc / ManNAc / HexNAc / glucose / mannose / hexose/ sugar 6-dehydrogenases were used for Case Study 1, and Lysozyme C and alpha-lactalbumin for Case Study 2. Sequences that were missed out by keyword search were obtained by iterative search i.e., searching Swiss-Prot using an HMM built from an initial set of sequences.

**Figure 1:**
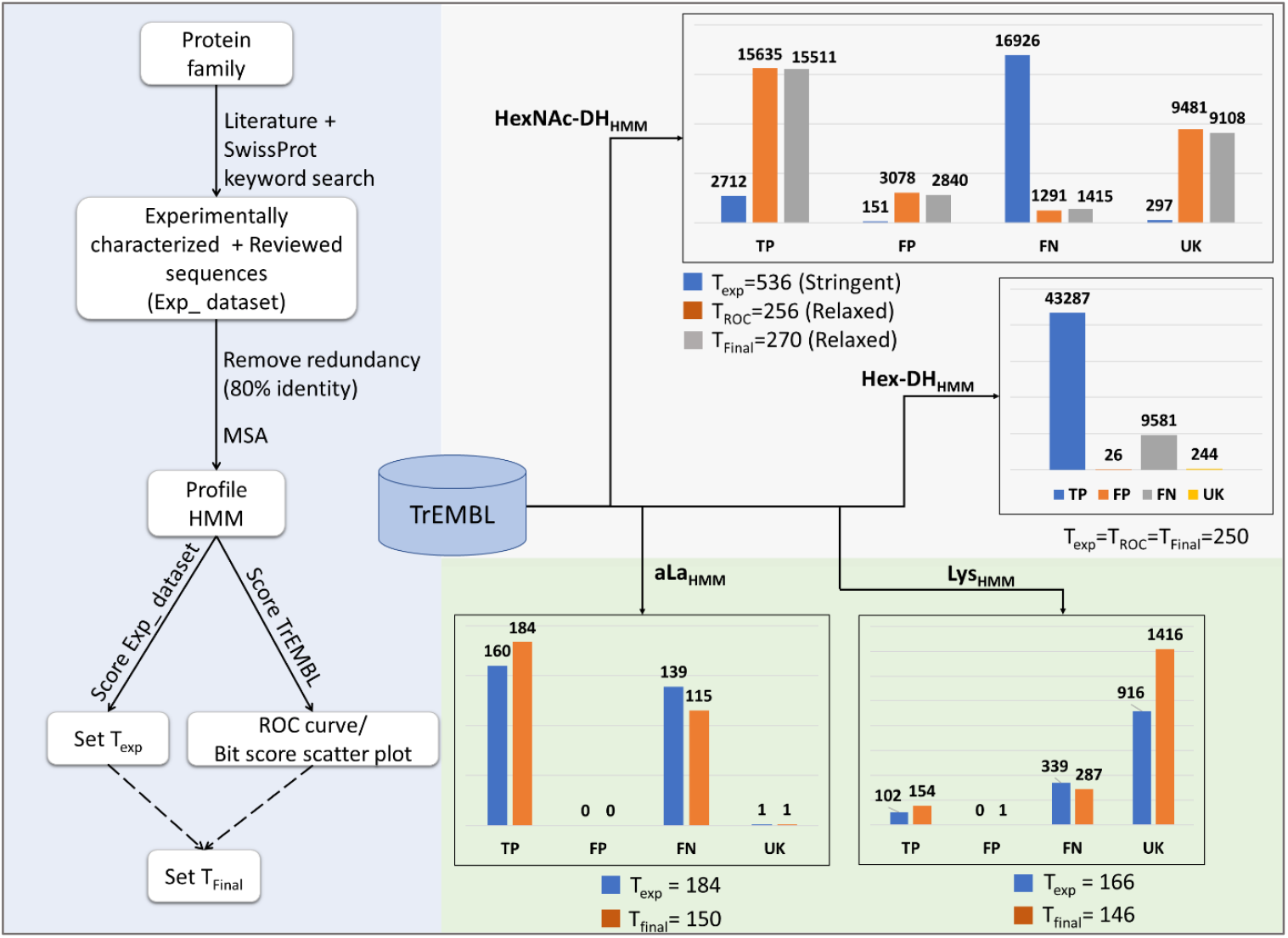
Procedure used to derive Profile HMMs and analyse hits for functional families in Case Studies 1 and 2. The final threshold, T_Final_, can be the same as T_exp_ or T_ROC_, or is set based on a bit score scatter plot. Details of Profile HMMs are given in Table 1. Details of proteins that constituted Exp_dataset are given in Supplementary_dataset.xlsx: NDP-6-DHs and lysozyme-C. TP and FN: True positives (bit score >= threshold) and false negatives (bit score < threshold) are proteins which have the same molecular function annotation as that assigned to the profile HMM. Here, threshold is T_exp_, T_ROC_ or T_Final_, as the case may be. FP: False positives (bit score < threshold) are proteins which have a molecular function annotation that is different from the one that is assigned to the profile HMM. UK: Proteins for which bit score >= threshold but there is incomplete or no molecular function annotation.

Bit scores were used as thresholds rather than E-values since they remain the same irrespective of the size of the database searched. Exp_dataset sequences were scored against the corresponding profile and the lowest bit score for Exp_dataset sequences was set as T_exp_. Entries in the TrEMBL database are automatically annotated using UniProtKB’s UniRule and Association-Rule-Based Annotator (ARBA) (UniProt Consortium 2021). These entries were used to generate Receiver Operator Characteristic (ROC) curves, as described in more detail elsewhere (Srivastava et al. 2020). In Case Studies 1 and 2, which involved functional families, hits to profiles were analysed for residue conservation using in-house python scripts. Such an analysis could not be done for Case Studies 3 and 4 since profiles in these Case Studies are meant to detect domains and domain boundaries rather than specific molecular functions.

**Table 1:**
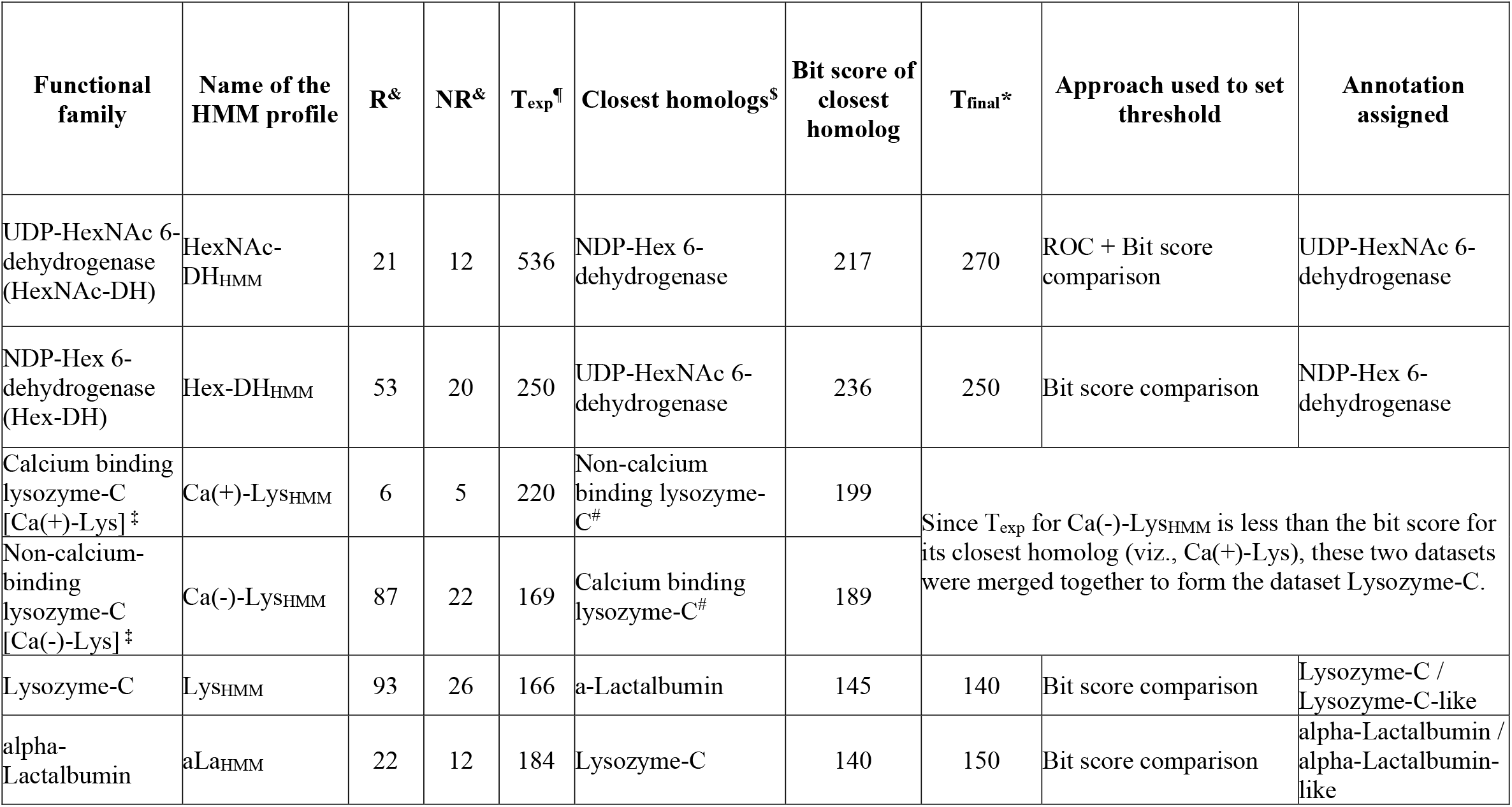
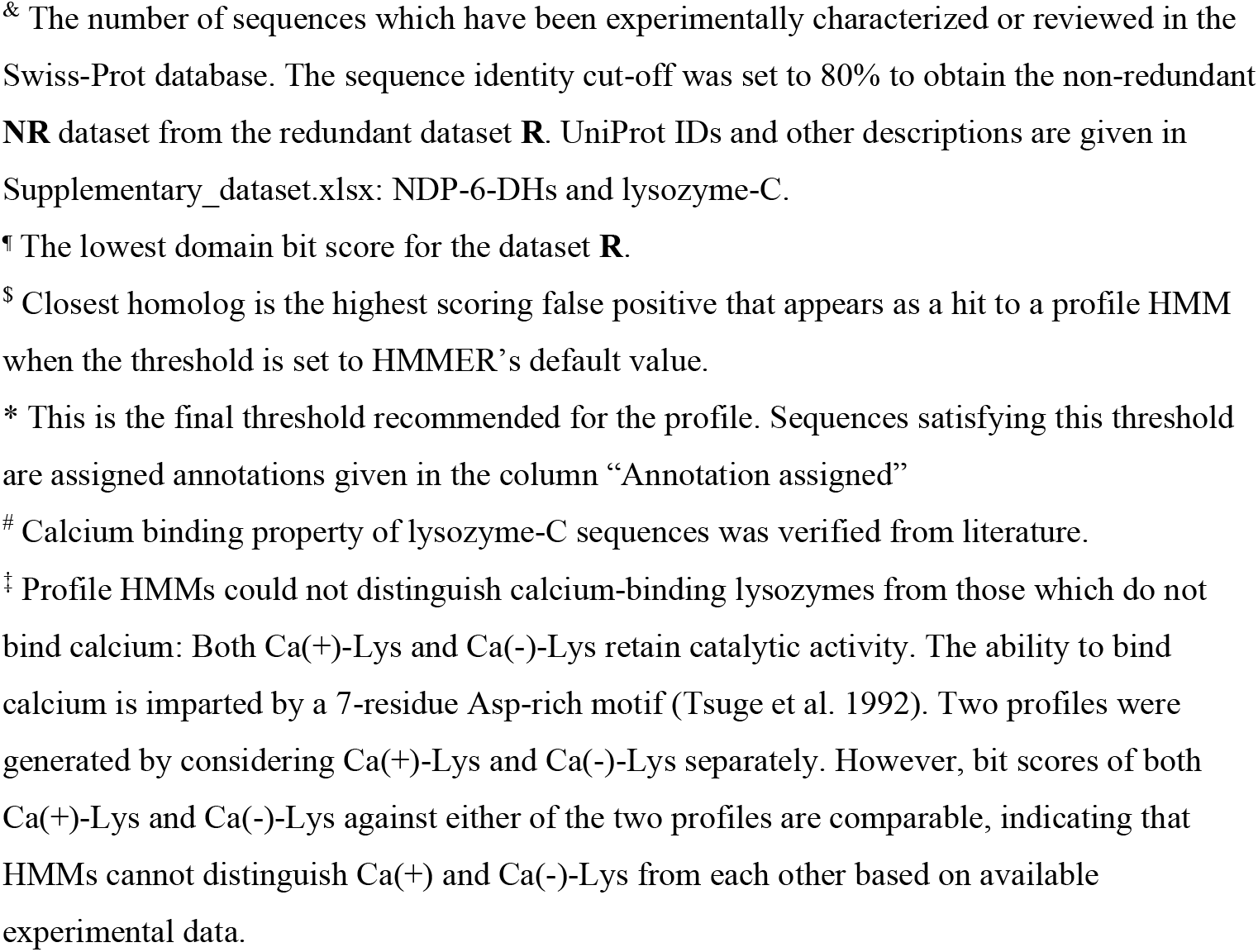
Details of profile HMMs built for Case Studies 1 and 2

To search for a sequence motif in a protein, position-specific amino acid frequency matrices were computed from MSA of the motif and converted into a position weight matrix of log odds score viz.,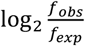, where *f*_*obs*_ and *f*_*exp*_ are observed and expected frequencies of occurrence of amino acid residues in a column of the MSA (i.e., a position in a motif). Expected frequencies were fetched from the TrEMBL database (Release 2021_01). For a given sequence, the *n*-residue stretch (*n* is the length of the motif) that has the highest log odds score was taken to be the best possible match for the motif; indeed, highest scoring motifs were subsequently verified to be the motifs of interest using in-house python scripts.

## RESULTS

### Case Study 1: Nucleotide sugar 6-dehydrogenase families

#### Generating Profile HMMs

Nucleotide sugar 6-dehydrogenases include HexNAc-DH and Hex-DH (Figure 2). They belong to the Short-chain Dehydrogenase Reductase (SDR) superfamily (Borg et al. 2020) and catalyze the conversion of hexose / HexNAc to the corresponding uronic acid. Two profiles, HexNAc-DH_HMM_ and Hex-DH_HMM_, were built to identify proteins belonging to these two functional families (Figure 1).

**Figure 2:**
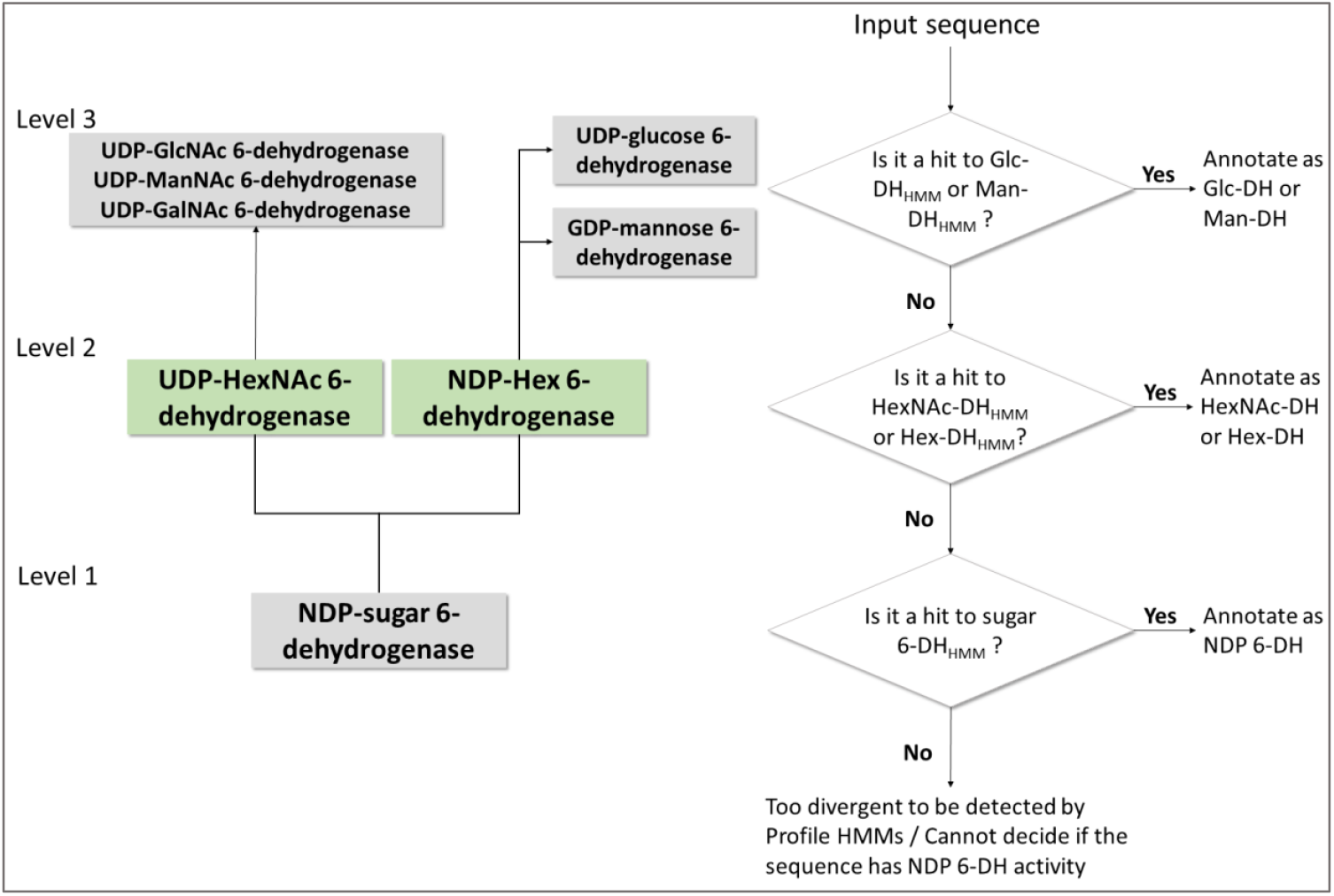
Categorization of nucleotide sugar 6-dehydrogenase family proteins into different levels based on substrate specificity. Assignment of substrate specificity is generic in Level 1 and increasingly specific in Levels 2 and 3. Separate profile HMMs could not be generated for the three HexNAc 6-DHs because of paucity of data and hence only a combined profile viz., HexNAc-DH_HMM_ could be generated. Rules on the right illustrate a stepwise approach to annotate an input sequence using profile HMMs built at three levels of categorization. Families whose profile HMMs are discussed in this study are shown in green boxes.

#### ROC curves

The TrEMBL database has nearly 17,000 sequences that are annotated as one of the HexNAc-DHs. Scoring TrEMBL using HexNAc-DH_HMM_ with T_exp_ (Table 1) as the threshold suggested that T_exp_ has poor sensitivity (Figure 1). To improve sensitivity, threshold was set to 256 bits (T_ROC_) using an ROC curve (Figure 3). With this threshold, there was a marked increase in sensitivity but with a concomitant increase in false positives (Figure 1). TrEMBL was scored using Hex-DH_HMM_ also with T_exp_ (Table 1) as the threshold. 81% of ∼53000 sequences annotated as one of the Hex-DHs in TrEMBL were obtained as hits along with a very small number of sequences as false positives; this implied that T_exp_ is optimal for Hex-DH_HMM_ (Figure 1). An ROC curve was generated to find a threshold with a higher sensitivity but for thresholds less than T_exp_, specificity decreased without much improvement in sensitivity (Figure 3).

**Figure 3:**
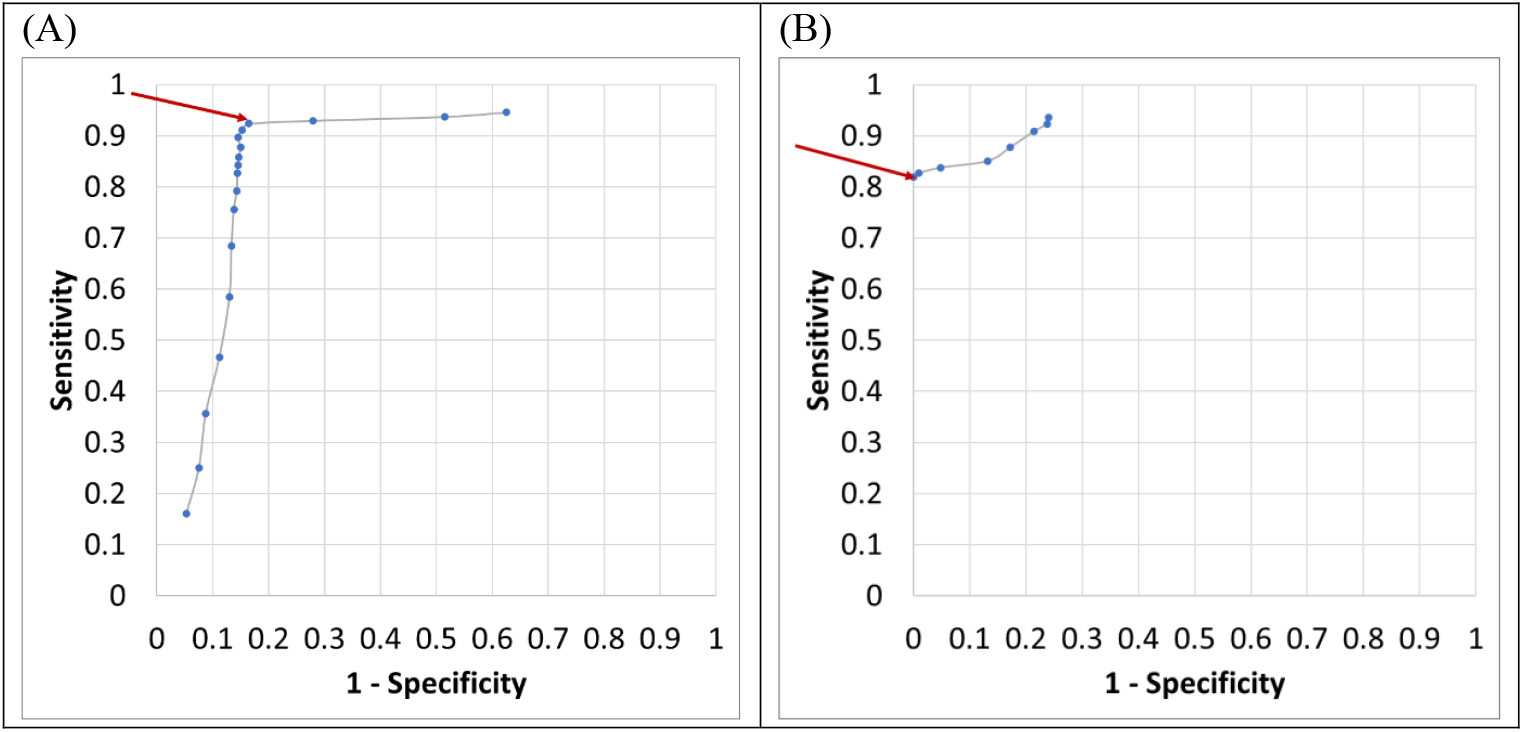
ROC curves for (A) HexNAc-DH_HMM_ and (B) Hex-DH_HMM_ obtained by scoring the TrEMBL database. Data points have been merely joined by a grey line. Arrow points to the data point correspond to the bit score threshold determined from the curve viz., T_ROC_.

#### Discriminating HexNAc-DH from Hex-DH

HexNAc-DH and Hex-DH are the closest homologs of each other (Table 1). 2800 out of 2840 false positives for the HexNAc-DH_HMM_ from TrEMBL are annotated as one of the Hex-DHs. To find-tune the threshold to improve specificity, TrEMBL was scored using both HexNAc-DH_HMM_ and Hex-DH_HMM_ using default values suggested by HMMER as thresholds. Plotting bit scores of hits common to the two profiles (Figure 4) showed that proteins which are “high-scoring” for one profile are “low-scoring” for the other. This plot showed that T_exp_ (250 bits) itself is optimal for Hex-DH_HMM_; however, T_Final_ for HexNAc-DH_HMM_ was revised marginally viz., from 256 to 270 bits (Table 1). Expectedly, this revision resulted in a marginal increase in specificity and decrease in sensitivity. Barring a few, false positives of HexNAc-DH_HMM_ score higher against HexNAc-DH_HMM_ as against Hex-DH_HMM._ Additionally, 98% of these false positives retain residues shown to be essential for catalysis in the HexNAc-DH family. Are these proteins actually HexNAc-DHs but are incorrectly annotated as Hex-DHs in TrEMBL?

**Figure 4:**
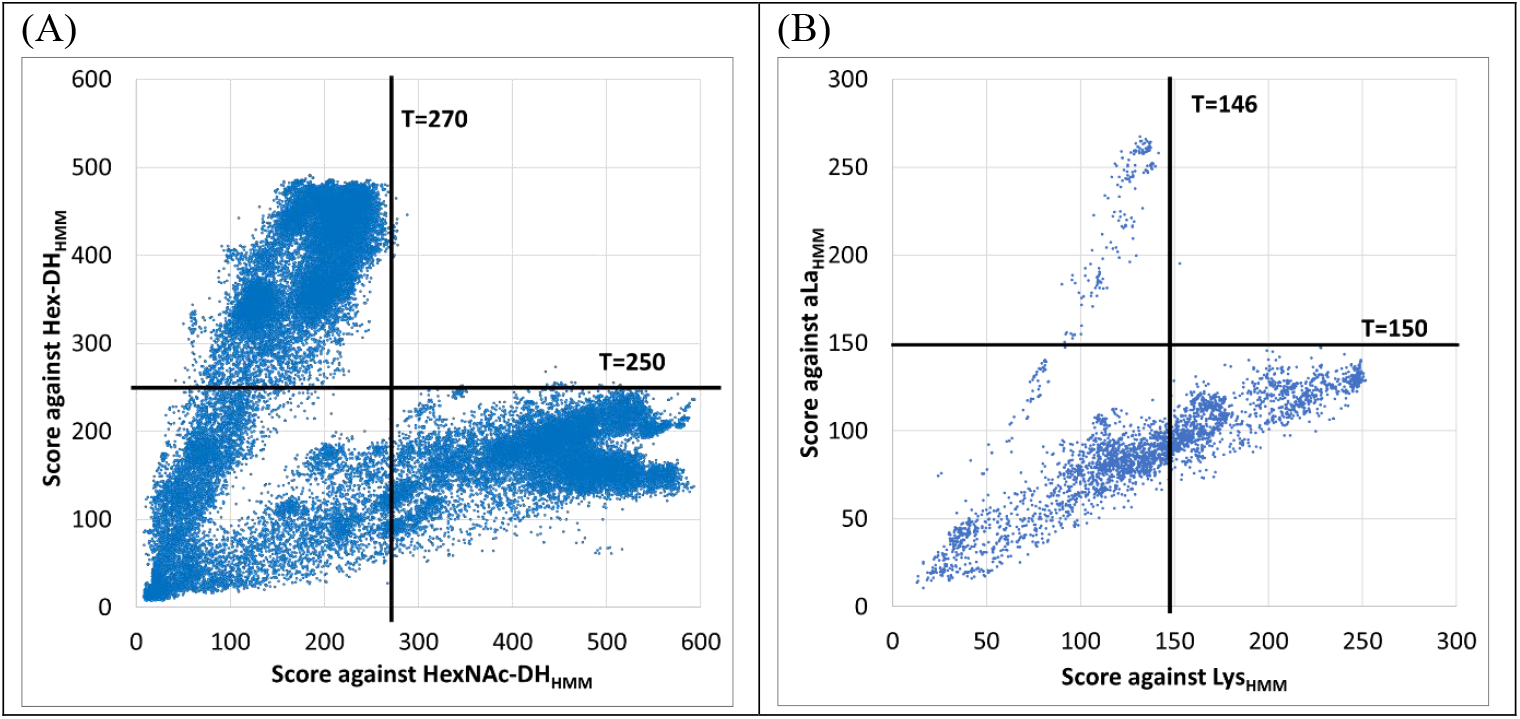
(A) Bit score comparison of TrEMBL entries that are hits to both Hex-DH_HMM_ and HexNAc-DH_HMM_. (B) Bit score comparison of common hits from TrEMBL against Lys_HMM_ and aLa_HMM_.

#### Residue conservation analysis

Conservation of active site residues among hits was examined to analyze the optimality of the threshold. In HexNAc-DHs, the cofactor binding motif GxGxxG, conserved cysteine, arginine and lysine (Lys204, Arg211 and Cys258, *Staphylococcus aureus* UDP-ManNAc 6-dehydrogenase numbering (Gruszczyk et al. 2011)) are implicated as important for catalysis. These are conserved in ∼96% of the sequences. Similarly, active site residues of NDP-hexose 6-dehydrogenases (Glu161, Lys210, Asn214, Lys218, Cys268 and Asp272, *Pseudomonas aeruginosa* GDP-mannose 6-dehydrogenase numbering) (Snook et al. 2003; Rocha et al. 2011) are conserved in ∼95% of the hits. Bit scores of sequences in which these residues are absent span the entire range. For instance, the second glycine of the GxGxxG motif is absent in 601 hits of HexNAc-DH_HMM_, whose bit scores range from 270-539. These observations indicate that profile HMMs are insensitive to the absence of critical residues i.e., the bit score can be higher than the threshold even when the mutation happens to be in a conserved region. Nevertheless, the number of false positives based on residue conservation are very low, suggesting that the threshold derived from aforementioned approaches is reliable. Thus,combining profile HMM with bit score distribution plot helped in discriminating families with different substrate specificities.

#### Fine-grained function annotation

HexNAc-DHs and Hex-DHs belong to the SDR superfamily and share the same fold (Gruszczyk et al. 2011; Rocha et al. 2011); none of the other functional families appear as hits to these profiles. Profile HMMs are able to capture sequence variations that give rise to utilization of either UDP-HexNAc or NDP-Hex. Using a similar approach as described above, we were also able to differentiate between UDP-glucose 6-dehydrogenases and GDP-mannose 6-dehydrogenases, enzymes that are strictly specific towards the respective substrates (Srivastava et al. 2020). However, it was challenging to resolve the substrate specificity among HexNAc-DHs because (i) some of these enzymes can utilize more than one substrate (Zhao et al. 2000) and (ii) experimental data on substrate specificity is available for very few HexNAc-DHs. Therefore, homologs of these enzymes identified using profile HMMs could, at best, be annotated as UDP-HexNAc 6-dehydrogenases. In scenarios where substrate specificity cannot be resolved using these profiles, an additional profile for all NDP 6-DHs can be used to capture homologs that are not hits to substrate-specific profiles. Using this set of profiles, rules for annotation can be formulated for an input sequence by scoring it at different levels, starting from the profile with highest substrate specificity, i.e., level 3 (Figure 2).

### Case Study 2: Lysozyme-C family

#### Sequence and structure homologs with mutually exclusive functions

Lysozyme-C family includes lysozymes, lysozyme-like proteins and alpha-lactalbumins (Prager and Wilson 1988; Grobler, Rao, et al. 1994; Irwin et al. 2011; Kalra et al. 2016). Some lysozymes bind calcium whereas others do not, denoted henceforth as Ca(+)-Lys and Ca(-)-Lys, respectively. alpha-Lactalbumins have lost catalytic activity but have acquired a new function wherein, as part of the lactose synthase complex, they modify the acceptor substrate specificity of beta-1,4-galactosyltransferase (Ramakrishnan and Qasba 2001). Phylogenetic analyses have indicated thatalpha-lactalbumins and lysozymes diverged from a common ancestor, a calcium binding protein (Prager and Wilson 1988; Irwin et al. 2011) (Figure S1).

Lysozyme-C family members exhibit high sequence and 3D structure similarities (Qasba and Kumar 1997) (Figure S2) despite of functional divergence. alpha-Lactalbumins (Prager and Wilson 1988) and lysozyme-like proteins (Irwin et al. 2011) share at least 30-40% sequence identity with lysozymes; this precludes the use of sequence identity alone for function annotation. Herein, we present a procedure to set bit score thresholds to distinguish alpha-lactalbumins from other members of this family.

#### Setting thresholds

Two profiles viz., Lys_HMM_ and aLa_HMM_ were generated for lysozyme and alpha-lactalbumins (Figure 1; Table 1). As mentioned earlier, T_exp_ led to poor sensitivity in the case of HexNAc-DH_HMM_, which meant that the threshold had to be lowered significantly. Is there a need for downward revision of T_exp_ of Lys_HMM_ and aLa_HMM_ also? It is noted that the highest score of an alpha-lactalbumin against Lys_HMM_ is 145 bits; similarly, the highest score of a lysozyme against aLa_HMM_ is 140 bits (Table 1). An ROC curve could not be plotted for Lys_HMM_ as there were no false positives scoring >145 bits among its hits from TrEMBL. Therefore, we used bit score comparison approach to set thresholds for both Lys_HMM_ and aLa_HMM_ (Figure 4). Accordingly, T_Final_ was set to 146 bits for Lys_HMM_ and 150 bits for aLa_HMM_ (Table 1). Case Studies 1 and 2 collectively show that a set of test sequences are required to adjust bit score threshold for profile HMMs that are built to annotate functions at fine grained level (Figure 2, Figure S1B).

#### Analysis of annotations of TrEMBL hits to Lys_HMM_

Using T_Final_ (Table 1) as the threshold, 1571 hits were obtained from TrEMBL. Of these, very few are annotated as lysozyme-C; many have incomplete or ambiguous annotations (“lysozyme”, “alpha-lactalbumin/lysozyme”, “lysozyme-like”) or are annotated as ‘sperm acrosome membrane-associated protein’, or do not have molecular function annotation (Figure 1). Several of these hits do not have the catalytic residues Glu53 and Asp70 (chicken egg-white lysozyme numbering (Malcolm et al. 1989)). Scores of such sequences fall in the range 146-236 bits. Sperm acrosome membrane-associate proteins are described in literature as lysozyme-like proteins, some of which have lost their bacteriolytic activity due to catalytic site mutations (Herrero et al. 2005; Wei et al. 2013: 6; Zhang et al. 2016). As observed earlier, profile HMMs are insensitive to point mutations and thus, catalytically active proteins could not be distinguished from inactive homologs using HMMs. To accommodate this limitation, the annotation ‘lysozyme-C / lysozyme-C-like’ was assigned for Lys_HMM_ (Table 1).

#### Analysis of annotations of TrEMBL hits to aLa_HMM_

Using T_Final_ (Table 1) as the threshold, 185 TrEMBL entries are obtained as hits, of which one has no molecular function annotation (Figure 1). 105 sequences are annotated as ‘Lactose synthase B’ and hence are true positives. Sequences of true positives were analysed for conservation of functionally important residues viz., Phe50, His51, Gln138 and Trp137 (bovine alpha-lactalbumin numbering (Grobler, Wang, et al. 1994)). Phe50 and His51 are conserved among all hits but Gln136 and Trp137, residues implicated in binding to galactosyltransferase (Grobler, Wang, et al. 1994), are absent in 55 sequences due to ∼20-30 residue C-terminal truncation. Such sequences score in the range 150-222 bits. The effect of such a truncation on lactose synthase activity is not known. Hence, the annotation ‘alpha-lactalbumin / alpha-actalbumin-like’ was assigned for the aLa_HMM_ (Table 1).

### Case Study 3: SH3 domain

#### A highly divergent sequence family

Src homology 3 (SH3) domains are ∼60-residue long structural elements characterized by a conserved beta-barrel fold (Figure S3) but no conserved sequence motif is found across all SH3 domains. Consequently, the Pfam database has 37 distinct families representing SH3 domains (Punta et al. 2012) (Supplementary_dataset.xlsx: SH3 families). SH3 domains are a part of multi-domain proteins that mediate diverse biological processes (Kurochkina and Guha 2013). Consequently, in databases such as TrEMBL, annotations for SH3 domain-containing proteins do not necessarily reflect the presence of this domain. Hence, unlike as in the case of single domain proteins, annotation-based approaches such as ROC curve are not suitable for defining bit score thresholds for SH3 domain profiles. Pfam assigns manually curated and periodically updated full-sequence and domain thresholds for every profile HMM, referred to as gathering thresholds by Pfam (Punta et al. 2012). Herein, we examine the domain gathering thresholds of SH3 domain profile HMMs (as of May 2021).

#### Are profile HMMs for all 37 families specific?

A pan-SH3 sequence dataset was created by pooling the seed alignment sequences of all 37 profiles. Scoring this dataset against all 37 profiles showed that some of the sequences satisfy the gathering threshold of more than one profile (Figure 5) implying that their gathering thresholds are less specific. Such overlaps, especially among families within a clan, are proposed to reflect evolutionary relationships between families (Punta et al. 2012). Nonetheless, they can be resolved by further optimization of thresholds or identification of additional approaches that can be used to distinguish between closely related families. Pfam does revise gathering threshold periodically (Punta et al. 2012).

**Figure 5:**
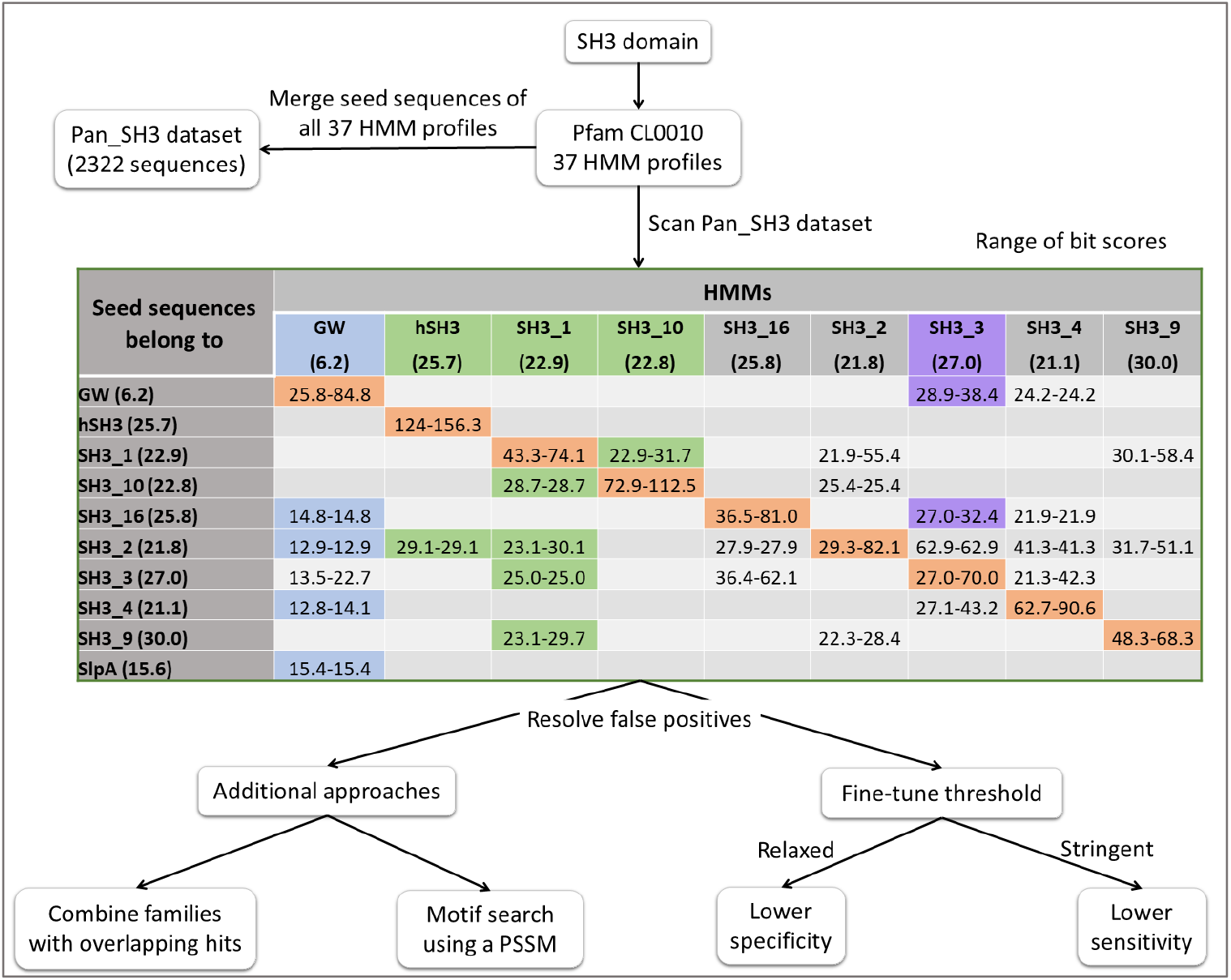
Steps used to analyse thresholds prescribed by Pfam for the 37 Profile HMMs for the SH3 domain. Seed sequences of all 37 profiles were scored against all 37 profiles. One or more seed sequences of 10 profiles (listed as rows) score above the domain gathering threshold of one or more profile (listed as columns) to which they do not belong (viz., false positives). Range of scores of seed sequences of a profile against own profile is highlighted in orange. Possible ways to resolve false positives are also depicted. Domain gathering thresholds of profiles are given in parenthesis following the name of the profile. Revising the gathering thresholds of hSH3 (from 25.7 bits to 30.0), SH3_1 (from 22.9 to 31) and SH3_10 (from 22.8 to 32 bits) excludes false positives (highlighted in green). For the profile GW (highlighted in blue), choosing the full sequence gathering threshold (21 bits) instead of the domain gathering threshold (6.2 bits) suffices to exclude most of the false positives. However, in the case of SH3_3 (highlighted in purple), a similar upward revision (from 27 to 38) leads to lower sensitivity concomitant with higher specificity. This holds true for SH3_16, SH3_2, SH3_4 and SH3_9 as well. The sequence Q5KV47_GEOKA/411-472 from the GW family scores 25.8 bits against GW profile but 38.4 and 24.2 bits against SH3_3 and SH3_4 profiles, respectively. The sequence coverage is higher against SH3_3 and SH3_4 profiles whereas it has long gaps in the seed alignment of GW family raising the possibility that is belongs to SH3_3 or SH3_4.

#### Revising thresholds of families with overlapping hits

In the case of the Gly-Trp (GW) domain profile, using the gathering threshold for the full sequence instead of that for the domain (https://pfam.xfam.org/family/PF13457#tabview=tab6) prevents sequence of SH3_2, SH3_4 and SH3_16 families from appearing as false positives to the GW profile (Figure 5). A similar upward revision of thresholds reduces false positives for SH3_1, hSH3 and SH3_10 also. However, such a revision compromises with the sensitivity of the profile in some cases (Figure 5). Under such a scenario, one may consider using two thresholds: (i) gathering threshold used by Pfam that has lower specificity (i.e., seed sequences of other SH3 families are hits) and assign a generic annotation viz., “SH3_1/hSH3/SH3_10/Gly-Trp domain” or (ii) use a revised, more specific threshold. For the remaining five families, multiple hits with scores comparable to that of seed sequences were obtained (Figure 5). To resolve false positives among hits of their profile HMMs, one can either employ residue/motif conservation analyses, or merge the seed sequences into a single profile and revise the annotation assigned to the profile (Figure 5). We will describe the motif conservation approach below.

#### Using family specific motifs to discriminate SH3 families

For this purpose, SH3_1 and SH3_2 families were considered as an illustrative example. Nearly 80% of SH3_1 seed sequences are hits to SH3_2 profile (Figure S4). Visual inspection of MSAs of seed sequences revealed a 6-residue extension in SH3_2 (Figure S5). To identify distinguishing motifs, SH3_1 and SH3_2 seed sequences were collectively submitted as input to MEME suite. Three motifs from the SH3_1 family are conserved in all the seed sequences, but their log-odds scores are inadequate to distinguish SH3_2 from SH3_1, either individually or in combination. Two motifs were identified in the SH3_2 family and these are conserved in 27 out of 28 seed sequences. One of these motifs is able to distinguish SH3_1 from SH3_2, with the exception of a false negative (the 28^th^ sequence in which motif is absent). Log-odds scores of seed sequences of both families obtained from the aforementioned five motifs are given as supplementary data (supplementary_dataset.xlsx: Log odds scores). Thus, by combining profile and motif search approach we can attain distinction between families that cannot be resolved based on profile HMMs alone.

### Case Study 4: GT-A fold of glycosyltransferases

#### GT-A fold

Glycosyltransferases (GTs) are enzymes that catalyse the transfer of sugar moieties from activated donor molecules to acceptor molecules forming glycosidic bonds (http://www.cazy.org/GlycosylTransferases.html). CAZy has nearly 0.83 million GTs which are grouped into 114 sequence-based families (as of May 2021) and GT2 is the largest of these families with ∼250,000 proteins. Most of the GTs whose 3D structure has been determined experimentally adopt either the GT-A or GT-B fold; few others have GT-C, GT-D or GT-E fold (Gloster 2014; Zhang et al. 2014; Kattke et al. 2019). CAZy has annotated that 19 GT families adopt GT-A fold including GT2. This is a similar scenario to that of SH3 family wherein 37 subfamilies are formed by Pfam based on sequence similarity, all of which adopt the SH3 fold.

#### Function annotation of multi-domain proteins

3D structures of several GT2 family members have been determined (Table S2). It can be seen from these studies that the lower limit for the number of amino acids in a domain that adopts the GT-A fold (henceforth referred to as the GT-A domain) is around 210 residues. There are ∼500 sequences in SwissProt that are labelled as GT2. The lengths of these proteins vary over a large range (Figure S6); this clearly suggests that many of them have one or more domains in addition to the GT-A domain. Functional annotation of such proteins necessarily involves the demarcation of domain boundaries, functional annotation of each domain as well as annotating the function performed by these covalently linked domains in unison. As the first step in this direction, since CAZy has assigned GT-A fold to GT2 family, a profile HMM-based approach was taken up to identify (i.e., find and demarcate domain boundary) GT-A domains in the GT2 family proteins (Figure 6).

**Figure 6:**
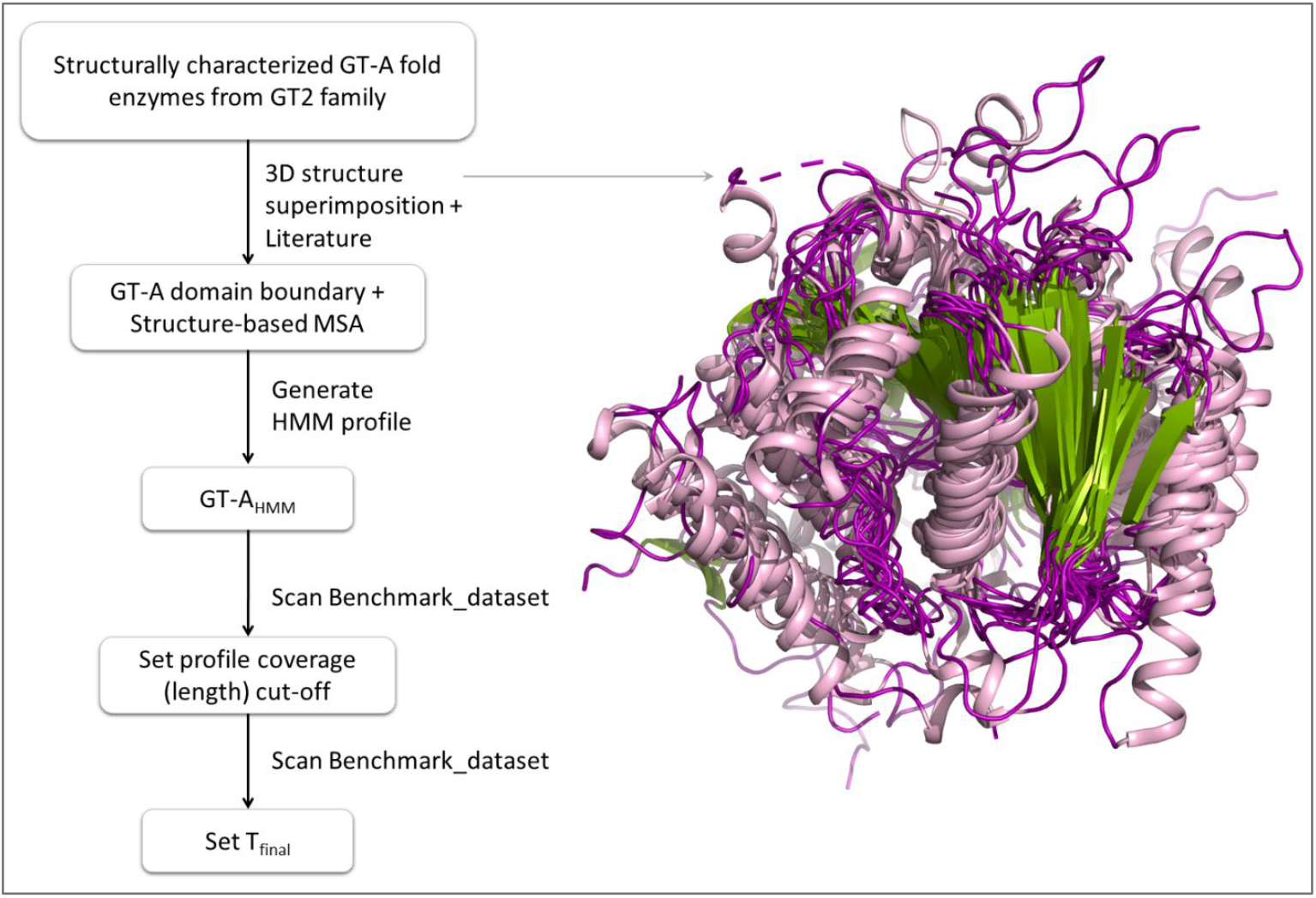
Steps followed to determine bit score threshold and HMM coverage cut-off for GT-A_HMM_. PDB Ids and related information for proteins used for superposition are given in Table S1, Supplementary_dataset.xlsx: GT2 structures.

#### MSA of GT-A fold domains

As mentioned earlier, 3D structures of 11 GT2 family proteins are known (Table S1). They vary in length from 256 to 978 amino acids, and thus, some are multi-domain proteins. GT-A domain boundaries in these proteins were identified from literature. Of these 11 proteins, chondroitin polymerase from *Escherichia coli* strain K4 (PDB id 2Z87) has two occurrences of the GT-A fold and these were taken as two separate GT-A domains. In all, the 12 GT-A domains were superimposed on each other using the Dali server (Figure 6). An MSA was derived from superimposed structures and this was used to generate GT-A_HMM_ (Figure 6). As has already been reported, MSAs derived from 3D structure superimposition are better than those derived solely from amino acid sequences (Carpentier and Chomilier 2019).

#### How to set bit score threshold for the GT-A_HMM_?

As in the case of SH3 domain containing proteins, GT-A domain containing proteins in the TrEMBL database are not expected to contain fold level annotations and hence using an ROC curve did not seem feasible. A curated dataset of sequences with experimental evidence (viz., enzyme activity assay, complementation studies, etc.) or those that have high sequence similarity to experimentally characterized sequences were chosen out of entries labelled as GT2 in SwissProt. This resulted in a benchmark dataset of 170 proteins (Supplementary_dataset.xlsx: GT2 benchmark dataset). The benchmark dataset was scored using GT-A_HMM_ with default threshold of HMMER. 137 sequences scored, whose domain bit score is in the range 3-172 with HMM coverage being as low as 15% (Supplementary_ dataset.xlsx: GT-A_HMM_ hits). However, bit score does not correlate with HMM coverage (Figure 7). HMM coverage of some of these sequences is <50 %, and the 33 sequences of the benchmark dataset do not score at all. Thus, it remains uncertain whether they adopt a GT-A fold. On the other hand, several hits have >70% coverage but score low, implying that they have diverged significantly while keeping the fold intact. These sequences do not match the C-terminal region of profile HMM (Supplementary_dataset.xlsx: GT-A_HMM_ hits). It is observed that from the MSA (Figure S7) that the C-terminal region (∼20 residues) of sequences used to build GT-A_HMM_ are less conserved. We therefore set 160 residues as the HMM coverage cut-off (∼78% of the length of the HMM profile). The lowest bit score of a sequence satisfying coverage cut-off is 25 bits, which was set as T_Final_ (Figure 7).

**Figure 7:**
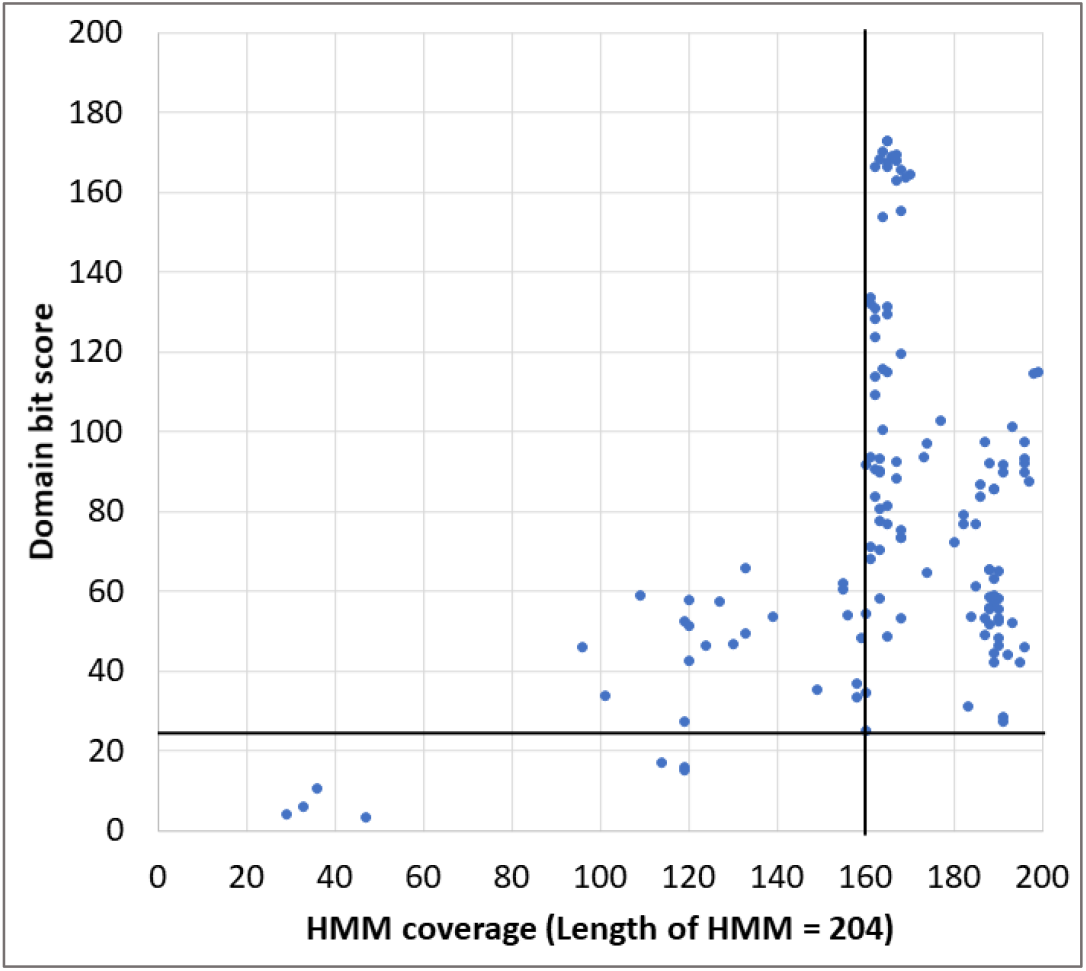
Bit scores of 133 sequences from GT2 benchmark dataset plotted against the corresponding HMM coverage (Supplementary_dataset.xlsx: GT-A_HMM_ hits). The horizontal line represents bit score cut off (=25 bits) and the vertical line represents HMM coverage cut-off (=160).

#### GT-A fold domain in other GT families of CAZy

GT-A_HMM_ was used to find domain homologs in other CAZy families that are annotated by CAZy as containing GT-A fold (GT6, GT7, GT8, GT15, GT27, GT64, GT81, etc.). Structurally characterized members of these GT families were scored using T_Final_. We found a sequence (UniProt accession number Q86SR1) belonging to GT27 satisfies both coverage cut-off and T_Final_ (Supplementary_dataset.xlsx: GT-A_HMM_ hits), suggesting that the GT-A domain of this 603 residue protein shares sequence similarity with the GT-A domain found in several GT2 family sequences. No other hits were obtained, implying that multiple profile HMMs are required to capture GT-A folds, as is the case with SH3 domains.

## Discussion

Profile HMMs are powerful tools to detect remote homologs in sequence databases and sequenced genomes. The default threshold set by HMMER search tool is an E-value of 10, which most often results in poor specificity. The false positives can be similar sequences that have diverged to perform different functions, or homologous fragments present as part of other polypeptides. Therefore, using profile HMMs for the identification of homologs necessitates manual intervention to set thresholds that can increase specificity. We have used four Case Studies to illustrate approaches that can be adopted to optimize bit score thresholds.

Case Studies 1 and 2 describe approaches to identify homologs of functional families. We set a threshold for the nucleotide sugar 6-dehydrogenase family using a combination of ROC curve and bit score comparison. The ROC curve approach relies on molecular function annotations of TrEMBL sequences. Assuming that the annotations are correct, ∼10% of the hits were found to be false positives against HexNAc-DH_HMM_ and almost all of these are annotated as Hex-DH. Almost all the false positives of HexNAc-DH_HMM_ scored more against HexNAc-DH_HMM_ than against Hex-DH_HMM_, and retain active site residues of HexNAc-DH. On the other hand, only a small fraction of hits (i.e., 26 out of 43,557 hits (Figure 1)) appeared as false positives against Hex-DH_HMM_. In the lysozyme-C family, we did not find any false positives against Lys_HMM_ and aLa_HMM_, but we did observe several hits of Lys_HMM_ missing or having incomplete annotations (Figure 1). Such a skew in false positive rate is a consequence of the bias in the TrEMBL database from the view point of the extent of representation of different functional families. Another reason could be that even though TrEMBL annotations are largely correct, some sequences are incorrectly annotated and this warrants curation of annotation using additional approaches such as bit score comparisons. Such a curation can also lead to assigning specific annotations to entries with incomplete or missing annotations as is exemplified in the case of HexNAc-DH family (Figure 1). We also learn that in situations where substrate specificity cannot be assigned due to paucity of data, e.g., ManNAc 6-dehydrogenases or GlcNAc 6-dehydrogenases, a combined profile with broad annotation, i.e., HexNAc-DH is the best that can be done.

Lys_HMM_ and aLa_HMM_ were generated from sequences belonging to Chordates (and Arthropods in case of Lys_HMM_) and hits of these profiles expectedly belong to the same phyla. HexNAc-DHs have not been identified in eukaryotes as yet, but Hex-DHs contain members from all domains. Nevertheless, hits of HexNAc-DH_HMM_ contain sequences from eukaryotes (plants, nematodes, arthropods) and virus, indicating that sequence divergence among HexNAc-DHs are taxa independent. However, this may not always be true. Experimentally characterized GDP-mannose pyrophosphorylases from eukaryotes and bacteria show more similarity within than across domains. In such cases, profile HMMs generated using sequences from only one domain (e.g., eukaryotes) will not be sensitive to sequences from the other domain.

How essential is the supplementation of ROC curve with a bit score distribution plot for arriving at optimal thresholds? If an HMM represents a functional family that is distinctly different from other functional families in its primary structure, then an ROC curve should suffice. This will be the case if we build an HMM to represent the NDP-sugar 6-dehydrogenase family (Figure 2). However, if one were to have two HMMs each of which represents functional families that vary from each other only at a few homologous positions in the primary structure, then supplementing ROC curves with a bit score distribution plot will significantly improve the optimality of thresholds. This is the case with HexNAc-DH_HMM_ and Hex-DH_HMM_, and with Lys_HMM_ and aLa_HMM_ (Table 1). The extent of sequence divergence that accounts for functional variation (substrate specificity in this case) is marginal necessitating the supplementation.

Case Studies 3 and 4 involve identifying homologs of SH3 and GT-A domains. Unlike as in Case Studies 1 and 2, ROC curve or any other existing annotation-based approaches cannot be utilized even for the identification of domain homologs as annotations of most of their hits do not include domain information. As there are no benchmark datasets to determine thresholds, there is a larger compromise with sensitivity. Pfam has subclassified Src-homology 3 (SH3) family into 37 subfamilies based on sequence similarity and assigned gathering thresholds (Punta et al. 2012). Scoring a dataset containing seed sequences of all the 37 subfamilies, we found overlapping hits for multiple profiles, suggesting the need for revisiting gathering thresholds. We find that thresholds for these families can be resolved by either (i) merging subfamilies that contain high scoring false positives from each other, (ii) using additional approaches such as motif search in combination with profile HMMs, or (iii) considering two thresholds, one with high specificity (stringent threshold) another with high sensitivity (relaxed threshold). There is no straightforward way to assign optimal thresholds and a compromise between specificity or sensitivity is inevitable in the absence of benchmark datasets. Unlike SH3 domains that are involved in protein-protein interactions, GT-A fold (Case Study 4) is responsible for catalytic activity. However, like SH3 domain, it is often found as a part of a multidomain protein. Often, sequence-based approaches transfer molecular function annotation of one domain to the entire protein (Purushotham et al. 2020). Under these scenarios, it is important to define domain boundaries as a part of functional annotation. We used structurally characterized glycosyltransferases from GT2 family (that adopt GT-A fold) to illustrate an approach to demarcate domain boundaries. A manually curated dataset of experimentally characterized sequences classified under GT2 family in CAZy (Lombard et al. 2014) was used as benchmark dataset to identify HMM coverage threshold. Additionally, sequences belonging to other CAZy families adopting GT-A fold were used to identify a bit-score threshold. Thus, a combination of HMM coverage and bit-score threshold are required to identify GT-A domains.

How to find a bit score threshold that is optimal for specificity even if the search is poorly sensitive? One approach is the use of “negative” datasets. For example, to set threshold for an HMM that represents a KEGG Ortholog (KO) family (considered as the positive dataset), KofamKOALA uses proteins belonging to other KO families as negative datasets (Aramaki et al. 2020). The distribution of bit scores for proteins belonging to the positive and negative datasets are used to arrive at the threshold. HMMERCTTER follows a similar approach and identifies thresholds for sequence clusters within a superfamily by comparing inter-cluster scores (Pagnuco et al. 2018). The advantage of these two methods is that they do not require subject matter experts (SMEs) and hence are scalable (i.e., amenable for automation). These methods work well for HMMs that represent sequence families that share broad molecular function e.g., nucleotide sugar 6-dehydrogenases (as at Level 1 in Figure 2). The approach discussed in the present study is effective even for relatively fine grained function annotation (as at Levels 2 and 3 in Figure 2) because it involves SMEs; obviously, this is not truly scalable.

## List of abbreviations

ARBA: Association-Rule-Based Annotator
BLAST: Basic Local Alignment Search Tool
GT: Glycosyltransferase
Hex-DH: NDP-hexose 6-dehydrogenases
HexNAc-DH: UDP-HexNAc 6-dehydrogenases
HMM: Hidden Markov Model
KO: KEGG Ortholog
MAFFT: Multiple Alignment using Fast Fourier Transform
MEME: Multiple Expectation maximizations for Motif Elicitation
MSA: Multiple Sequence Alignment
NDP: 6-DH Nucleotide sugar 6-dehydrogenase
NDP-Hex: NDP-hexose
PSI-BLAST: Position-specific iterative
BLAST: ROC Receiver Operating Characteristic
UDP-HexNAc: UDP-N-acetyl hexosamine

## AUTHOR CONTRIBUTIONS

PVB conceptualized research, JS performed analysis, RM contributed to lysozyme-C and GT2 family analysis, AK contributed to SH3 domains analysis, PVB and JS cowrote the paper.

## ACKNOWLEDGEMENTS

We thank Toshi Mishra and Busi Rakesh for useful discussions. Jaya Srivastava is thankful to the Council of Scientific and Industrial Research, Government of India, for a research fellowship [number 09/087/(0877)/2017-EMR-I].

