## Supplementary_material for "Searching sequence databases for functional homologs using profile HMMs: how to set bit score thresholds?"

Powai

Mumbai 400076

India.

**Contents**

| **Data** | **Description** |
| --- | --- |
| Figure S1 | Composite schematic showing A) the phylogenetic relationship among proteins of lysozyme-C family and B) functional hierarchy in lysozyme-C family |
| Figure S2 | Structural similarity between lysozyme-C and alpha-lactalbumin |
| Figure S3 | 3D structure of a SH3 domain |
| Figure S4 | Bit score range of seed sequences of certain SH3 profiles against certain other SH3 profiles. |
| Figure S5 | MSA of seed sequences of SH3_1 and SH3_2 family |
| Figure S6 | Length variations in reviewed sequences belonging to GT2 family |
| Figure S7 | Structure based sequence alignment of sequences used to build GT-A_HMM_ |
| Table S1 | List of software and web servers used in the present study |
| Table S2 | Description of structurally characterized enzymes of GT2 family |
| References | References cited in supplementary figures |

**MS-EXCEL file provided separately:** Supplementary_dataset.xlsx

| NDP 6-DHs | Details of HexNAc-DH and Hex-DH sequences used to generate profiles for Case Study 1 |
| --- | --- |
| Lysozyme-C | Details of lysozyme-C family sequences used to generate profiles for Case Study 2 |
| SH3 domain | Details of 37 profile HMMs obtained from Pfam for the SH3 domain |
| Log odds Scores | PSSM log odds scores of seed sequences of SH3_1 and SH3_2 profiles |
| GT2 structures | GT2 family proteins with known 3D structure. These structures were superimposed on each other, an MSA for the region corresponding to the GT-A fold was derived from this superposition and this, in turn, was used to generate the HMM profile GT-A_HMM_ |
| GT2 benchmark dataset | Curated dataset of reviewed GT2 family sequences obtained from SwissProt |
| GT-A_HMM_ hits | *hmmsearch* output of hits to GT-A_HMM_ |

(A)

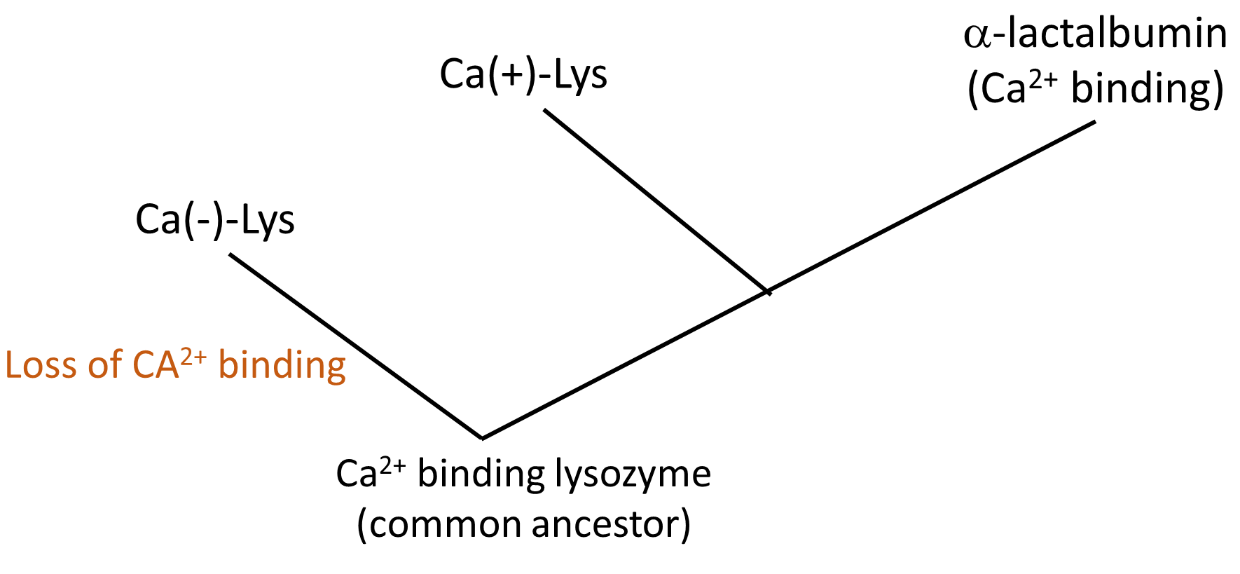

(B)

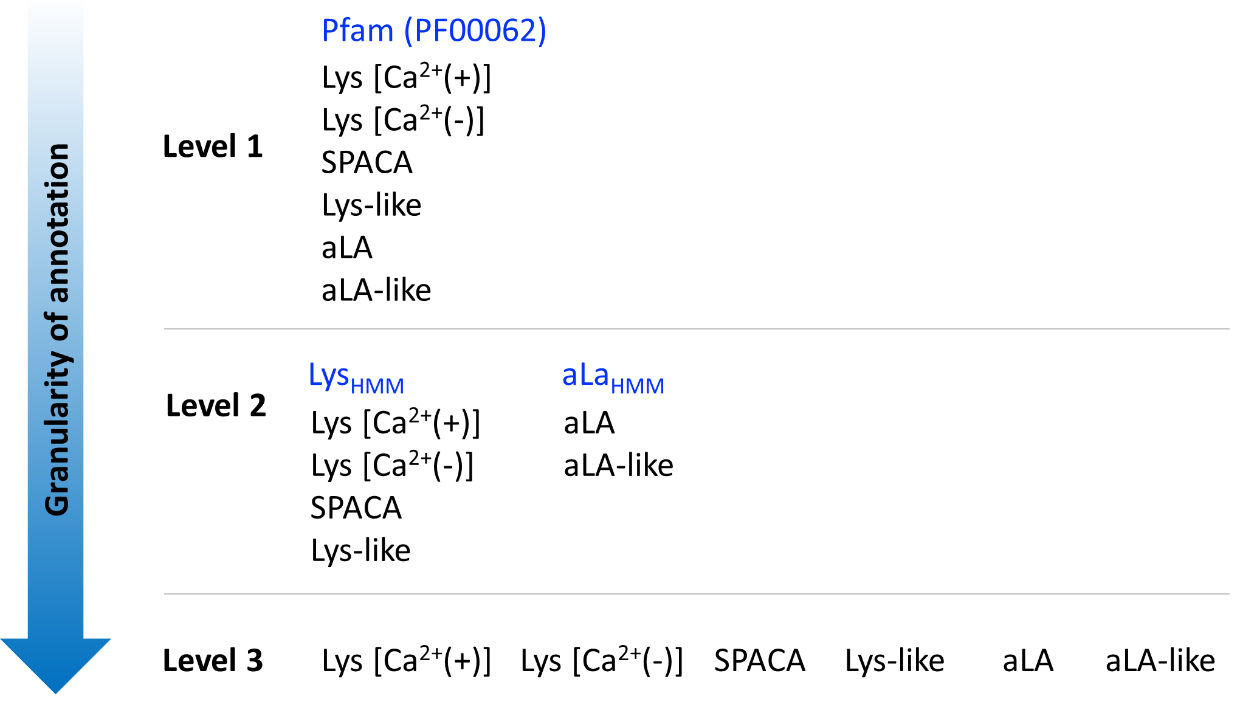

**Figure S1**: (A) A schematic showing the phylogenetic relationship between alpha-lactalbumin, Ca(+)-Lys and Ca(-)-Lys. Existing literature suggests that the common ancestor could bind Ca^2+^ (Grobler et al. 1994). Lysozyme-like proteins also belong to this family but data about their divergence is inadequate.

(B) Categorization of Lysozyme-C family proteins into different levels based on functional heterogeneity. Profile HMMs for level 1 are available on Pfam, for level 2 are generated in this study, and could not be generated for level 3 due to inadequate experimental data. SPACA and Lys-like proteins are expressed in mammalian testis and epididymis, and may or may not have antibacterial activity (Zhang et al. 2005; Narmadha et al. 2011: 4; Wei et al. 2013: 6). Even though Ca(+)-Lys and alpha-lactalbumin are suggested to diverge from a common ancestor, Ca(+)-Lys are more similar in sequence to Ca(-)-Lys. Note: Lys-like proteins mentioned here and in literature are different from Lysozyme_like Pfam family (PF13702). The latter bear no sequence similarity to members of lysozyme-C family.

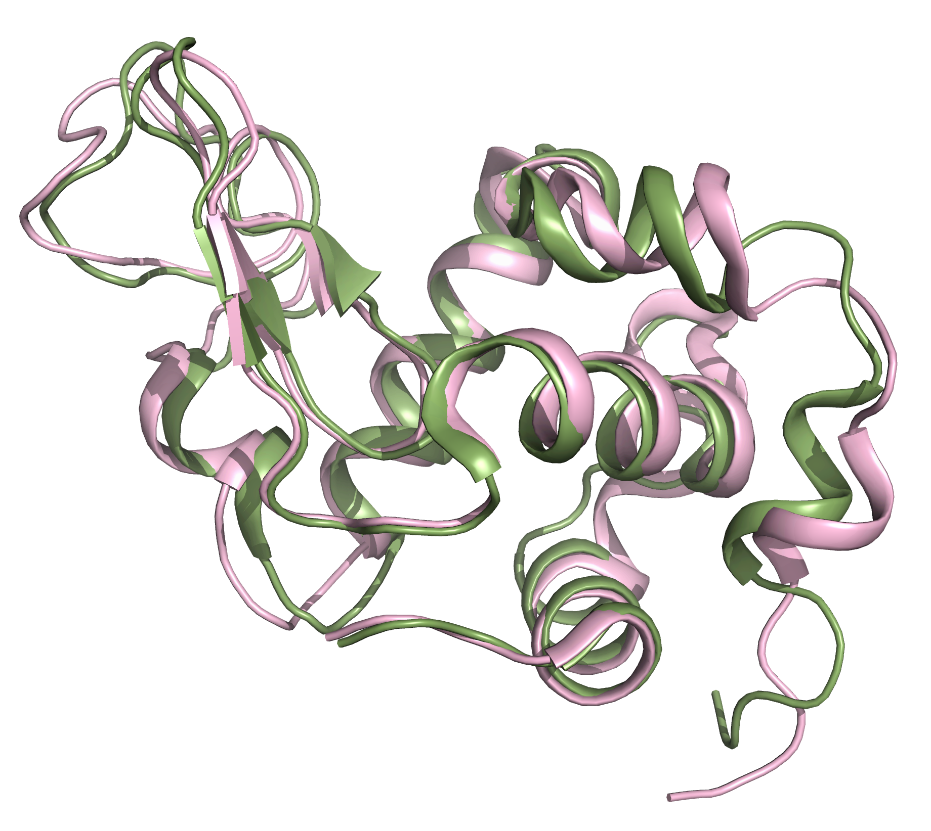

**Figure S2**: Structural superimposition of lysozyme-C from *Gallus gallus* (PDB: 2VB1, green) and alpha-lactalbumin from *Homo sapiens* (PDB: 1HML, pink) showing the high level of fold conservation.

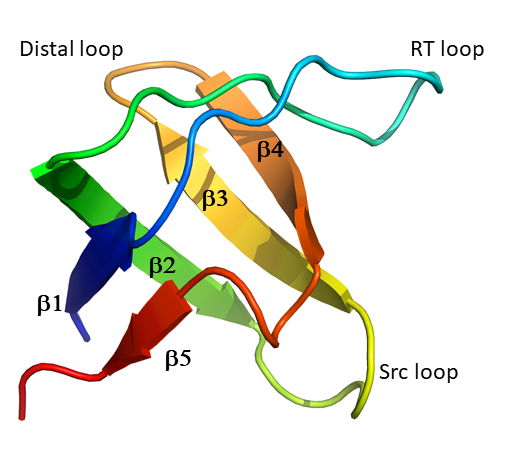

**Figure S3**: Rendering of the 3D structure of a SH3 domain (PDB ID: 1SHG). This domain is part of the chicken cytoskeletal protein spectrin. Strand and loop nomenclatures are as in (Mayer and Eck 1995).

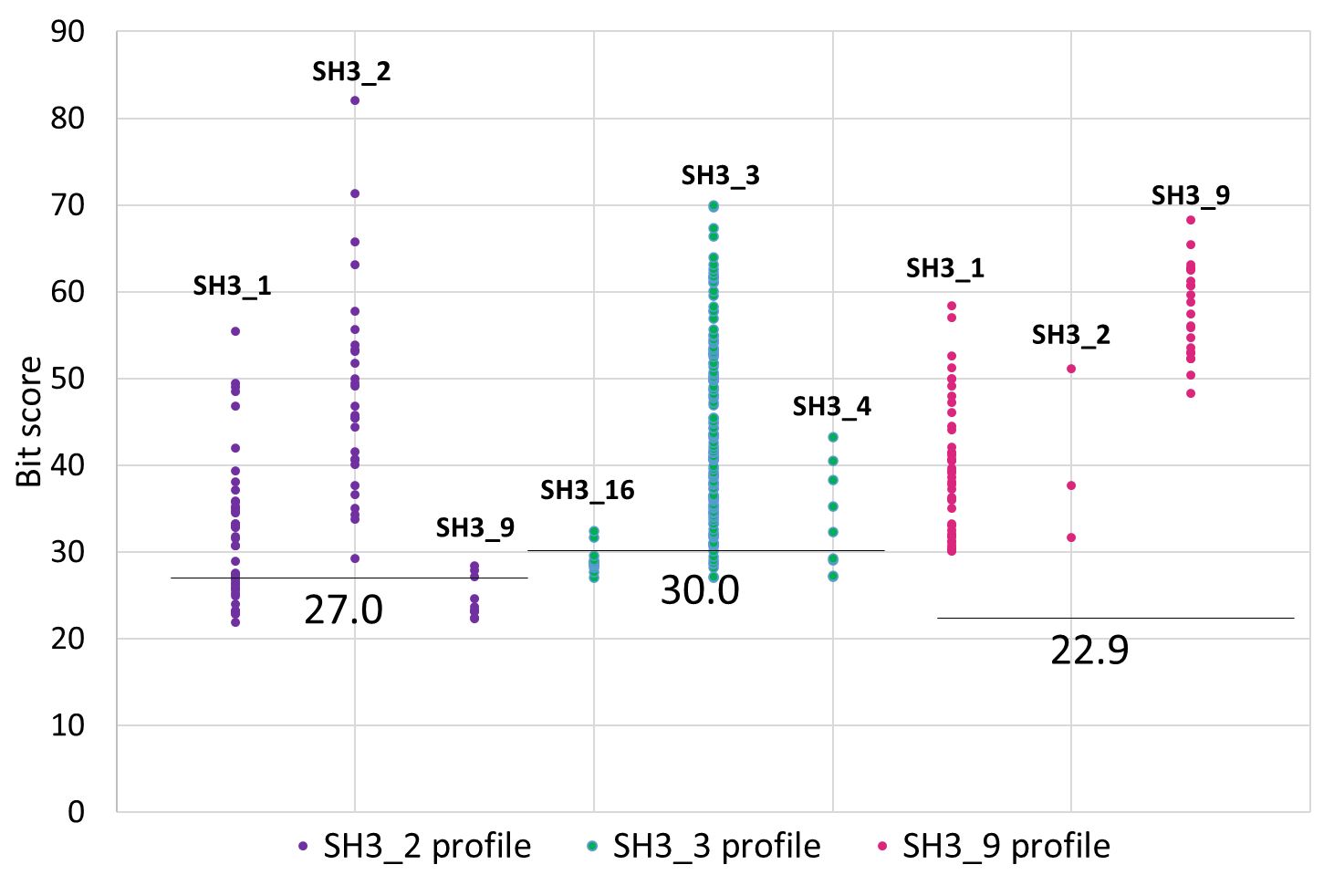

**Figure S4**: Bit score range of seed sequences of indicated HMMs against the SH3_2 (purple), SH3_3 (teal) and SH3_9 (pink) profiles. Gathering thresholds are mentioned just below the corresponding horizontal lines.

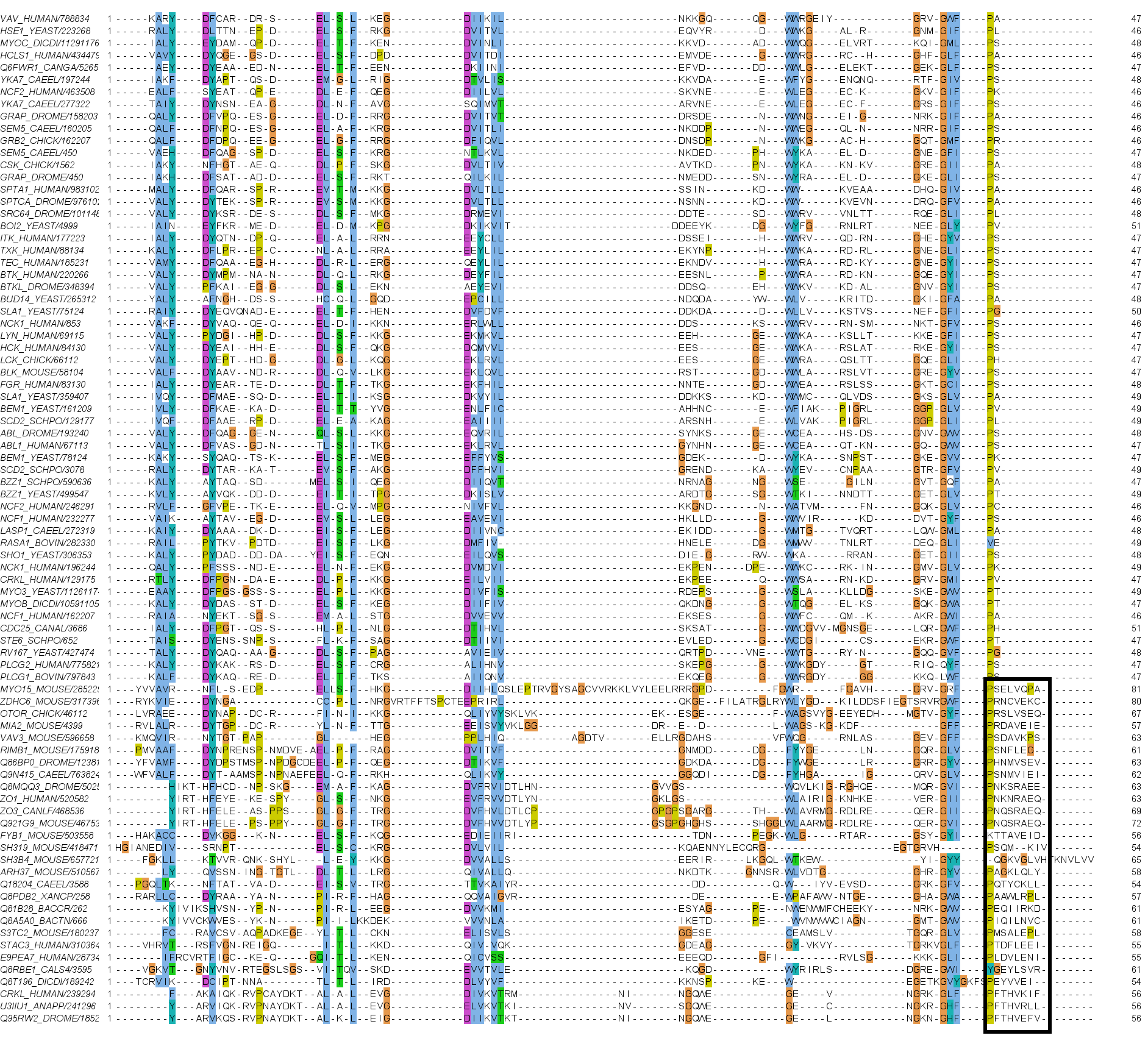

**Figure S5:** MSA of seed sequences of SH3_1 (55 sequences, top) and SH3_2 (28 sequences, bottom). Six amino acid extension in SH3_2 family is shown in rectangular box.

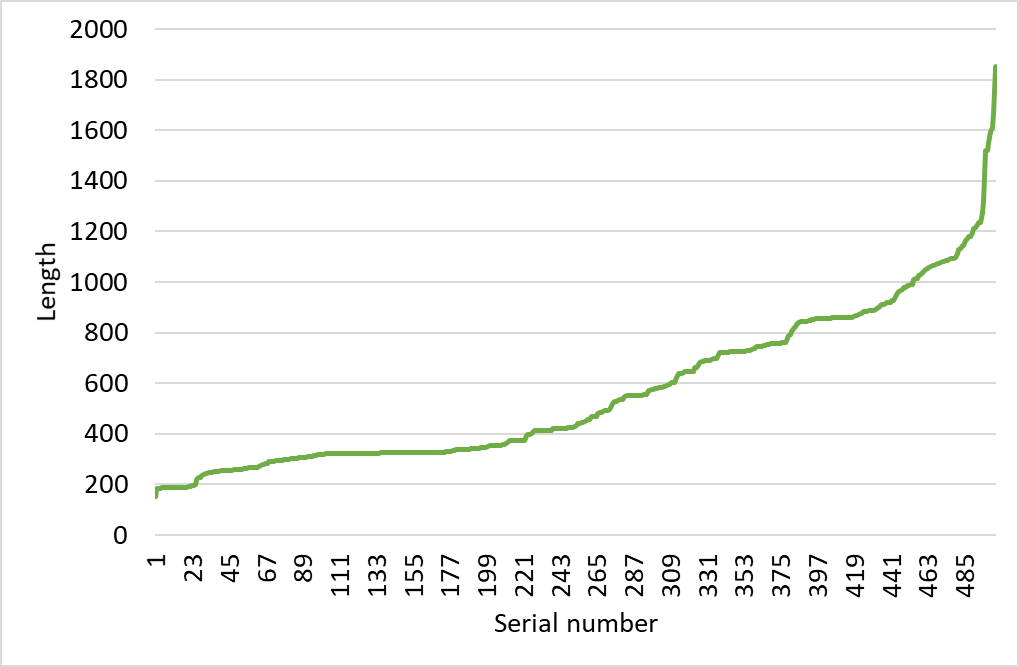

**Figure S6**: Lengths of the 504 structurally characterized or reviewed (Swiss-Prot) GT2 sequences. length is expressed as the number of amino acid residues in the polypeptide chain. Proteins were sorted in the increasing order of lengths merely for the purpose of making this plot.

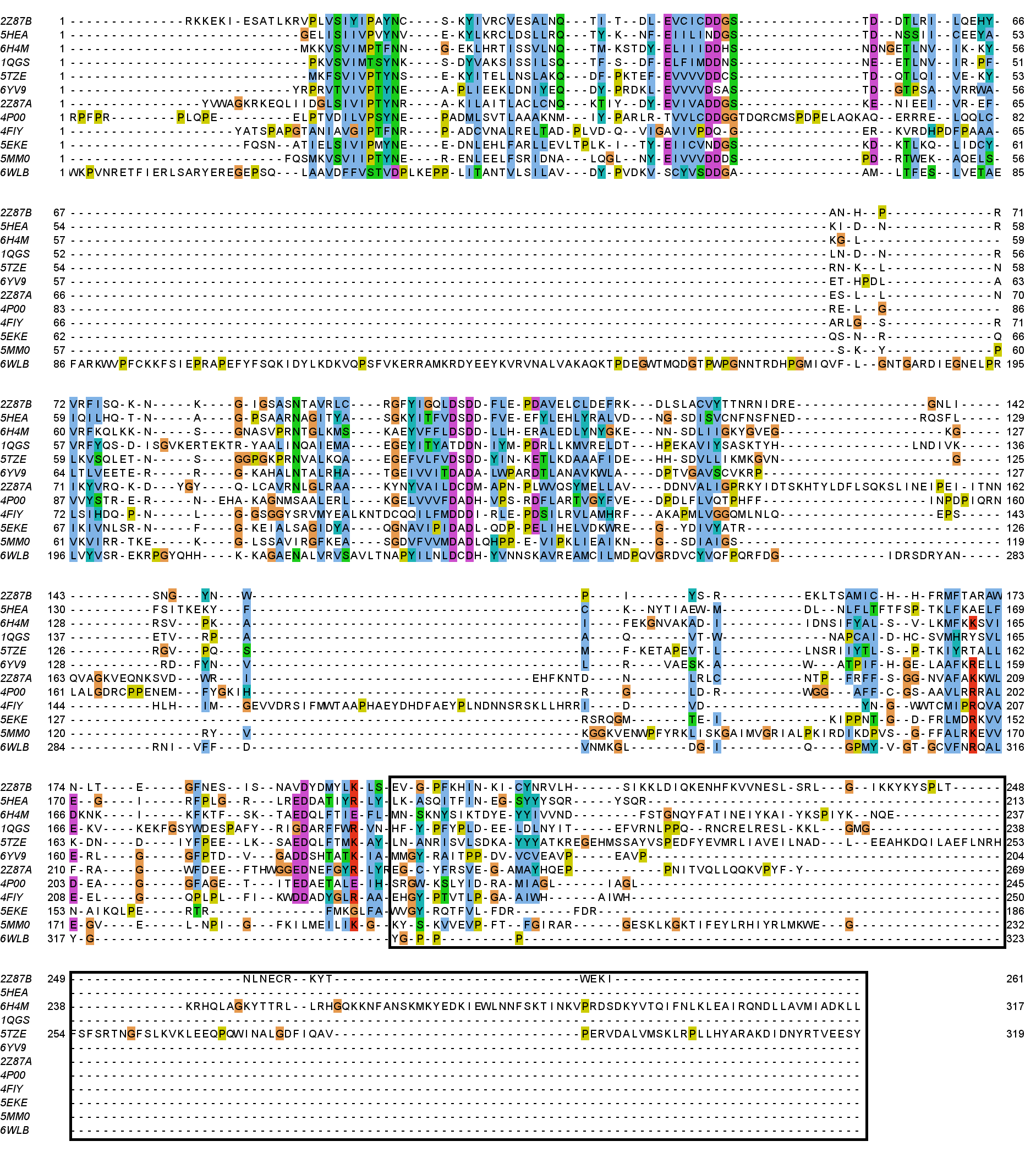

**Figure S7**: Structure based sequence alignment of GT2 family sequences (Table S1, Supplementary_dataset.xlsx: GT2 structures). The variable region at the C-terminus is shown in rectangular boxes.

**Table S1:** List of software and web servers used in the present study^¶^

| **Software / web server (version)** | **Purpose** | **URL (Reference)** |
| --- | --- | --- |
| UniProtKB | Amino acid sequences | (UniProt Consortium 2019) |
| Protein Data Bank | 3D structures | (Berman et al. 2000) |
| CD-Hit^§^ | Generating a non-redundant dataset | (Huang et al. 2010) |
| BLAST^‡^ | Pairwise sequence comparison | (Altschul et al. 1990) |
| PyMol | 3D structure visualization and comparison | (Schrödinger, 2021) |
| MAFFT^†^ | Multiple sequence alignment | (Katoh and Standley 2013) |
| Dali server | Structure-based sequence alignment | (Holm and Laakso 2016) |
| HMMER  *hmmbuild*  *Hmmsearch* | Build profile HMMs  Search sequence databases | (Finn et al. 2011) |
| CAZy | GT2 family sequences | (Lombard et al. 2014) |
| Pfam | Profile HMMs for SH3 domains | (El-Gebali et al. 2019) |
| STREME | Sequence motifs | (Bailey 2021 Mar 24) |

^¶^ All software and web servers were used with default values for all the parameters unless stated otherwise.

^§^ Sequence identity cut-off was set to 80%. The choice of the cut-off was primarily dictated by the paucity of experimental data.

^‡^ Both locally installed and online servers were used.

^†^ Command line application

**Table S2**: GT-A domain-containing GT2 family members^¶^

| **PDB ID** | **Organism** | **Molecular function** | **Protein length (a.a.)** | **GT-A domain boundary** | **GT-A domain length (a.a.)** |
| --- | --- | --- | --- | --- | --- |
| 2Z87 | *Escherichia coli* | beta 1,4-GalNAc transferase | 686 | 130-417 | 288 |
|  |  | beta 1,4-GlcA transferase |  | 418-682 | 265 |
| 5HEA | *Streptococcus parasanguinis* | GlcNAc transferase | 582 | 286-582 | 297 |
| 6H4M | *Staphylococcus aureus* | GlcNAc transferase | 327 | 1-327 | 327 |
| 1QGS | *Bacillus subtilis* | Glycosyltransferase | 256 | 1-256 | 256 |
| 5TZE | *Staphylococcus aureus* | GlcNAc transferase | 573 | 1-319 | 319 |
| 6YV9 | *Pyrobaculum calidifontis* | Dolichol phosphate mannosyltransferase | 365 | 41-250 | 212 |
| 4P00 | *Rhodobacter sphaeroides* | Beta-1,4 glucosyltransferase | 788 | 128-368 | 241 |
| 4FIY | *Mycobacterium tuberculosis* | Processive galactofuranosyltransferase with alternating β-1,5 and β-1,6 linkages | 637 | 151-396 | 246 |
| 5MM0 | *Pyrococcus furiosus* | Dolichol phosphate mannosyltransferase | 352 | 1-229 | 229 |
| 5EKE | *Synechocystis* sp. | Polyisoprenyl-glycosyltransferase | 318 | 1-206 | 206 |
| 6WLB | *Populus tremula* | Beta-1,4 glucosyltransferase | 978 | 225-546 | 322 |

^¶^ UniProtKB accession number and PubMed IDs are given in Supplementary_dataset.xlsx: GT2 structures.
